# Palmitoylethanolamide shows limited efficacy in controlling cerebral cryptococcosis *in vivo*

**DOI:** 10.1101/2023.04.10.536237

**Authors:** Melissa E. Munzen, Marta Reguera-Gomez, Mohamed F. Hamed, Vanessa Enriquez, Claudia L. Charles-Nino, Michael R. Dores, Karina Alviña, Luis R. Martinez

## Abstract

*Cryptococcus neoformans* (*Cn*) is an encapsulated neurotropic fungal pathogen and the causative agent of cryptococcal meningoencephalitis (CME) in humans. Recommended treatment for CME is Amphotericin B (AmpB) and 5-fluorocytosine (5-FC). Though effective, AmpB has displayed numerous adverse side effects due to its potency and nephrotoxicity, prompting investigation into alternative treatments. Palmitoylethanolamide (PEA) is an immunomodulatory compound capable of promoting neuroprotection and reducing inflammation. To investigate the efficacy of PEA as a therapeutic alternative for CME, we intracerebrally infected mice with *Cn* and treated them with PEA or AmpB alone or in combination. Our results demonstrate that PEA alone does not significantly prolong survival nor reduce fungal burden, but when combined with AmpB, PEA exerts an additive effect and promotes both survivability and fungal clearance. However, we compared this combination to traditional AmpB and 5-FC treatment in a survivability study and observed lower efficacy. Overall, our study revealed that PEA alone is not effective as an antifungal agent in the treatment of CME. Importantly, we describe the therapeutic capability of PEA in the context of *Cn* infection and show that its immunomodulatory properties may confer limited protection when combined with an effective fungicidal agent.

## INTRODUCTION

*Cryptococcus neoformans* (*Cn*) is an encapsulated neurotropic fungal pathogen and the causative agent of cryptococcosis in humans. *Cn* causes opportunistic disease in HIV-infected individuals due to low counts of CD4^+^ T cells, with this mycosis accounting for approximately 15 to 19% of AIDS-related deaths (1, 2). Cryptococcal meningoencephalitis (CME) is the most severe manifestation of the disease and results in an estimated 181,100 deaths annually worldwide. CME is a major cause of death in sub-Saharan Africa, where 2.5 million individuals are positive for HIV, over 58% of the global HIV-positive population (3). In fact, sub-Saharan Africa carries an estimated 73% of the global deaths related to CME annually (2). Despite the efficacy of and increasing accessibility to antifungal therapy, the high incidence of death from cryptococcal infection is associated with low adherence to antiretroviral therapy (4), as well as the lack of licensed and available treatment options (5).

The World Health Organization places cryptococcosis as a high priority mycosis and recommends a combination of amphotericin B (AmpB) and 5-flurocytosine (5-FC), followed by fluconazole as the therapeutic management of CME in severely immunosuppressed individuals (6). There are limitations to this treatment regimen, particularly with AmpB being highly toxic causing nephrotoxicity, phlebitis, hypokalaemia, and anaemia (7), often requiring hospitalization (8), and the high costs and limited accessibility of 5-FC (9). Various formulations of AmpB have been tested, such as single-dose liposomal AmpB (10), which has shown similar efficacy to conventional AmpB with fewer complications, and AmpB colloidal dispersion (11). Different combinations of antifungal therapies with AmpB have also been reviewed, such as tamoxifen (12), but fungal clearance and recovery rates have not been as effective.

Palmitoylethanolamide (PEA) is an endogenous lipid compound capable of controlling and reducing neuroinflammation (13, 14). PEA downregulates the expression of inducible nitric oxide synthase and NF-κB nuclear translocation through interaction with cellular receptors, notably the peroxisome-proliferator-activated receptor alpha, present in many cell types including microglia (15). Modulation of microglial phenotypes to promote phagocytosis and migration have been observed with PEA treatment (16, 17), as well as anti-inflammatory effects, neuroprotection, reduced apoptosis, and prolonged neuronal survival (18–20). PEA enhances survival in mice infected with *Escherichia coli* by modulating inflammation and stimulating bacterial phagocytosis by microglia (21). Interestingly, PEA also acts on several molecular targets both in the central and peripheral nervous systems, making it an attractive therapeutic agent for many central nervous system-related diseases (22). In fact, a clinical trial in the 1970s in the former Czechoslovakia involving 3,600 patients demonstrated that PEA had no adverse effects at daily doses of 600 to 1,800 mg (23, 24).

The incidence of CME and limited availability of accessible, safe, and effective treatments warrant further investigation into other potential therapeutic agents. Hence, in this study, we investigated the efficacy of PEA alone or combined with other commonly used antifungal drugs (e.g., AmpB and 5-FC) during *Cn* brain infection. Using a stereotaxic model of intracerebral (i.c.) mouse infection we evaluated the potential of PEA as a candidate for the clinical management of CME.

## RESULTS

### AmpB and PEA combination extend the survival of C57BL/6 mice i.c.-infected with *Cn*

We first assessed the impact of PEA alone or combined with AmpB in combating cerebral cryptococcosis (Fig. 1). Using a stereotaxic apparatus (Fig. 1A), we i.c.-infected 10^4^ *Cn* strain H99 cells into the mouse striatum (*n* = 7 mice per group, 6-8 weeks old, female), following the brain coordinates described previously (25). The striatum is a region in the basal ganglia of the brain that coordinates multiple aspects of cognition and motor function. We used it as the site of infection because this brain region is frequently affected in patients with CME (26) with cryptococcoma formation around the Virchow-Robin spaces (i.e., perivascular fluid-filled cavities that surround perforating blood vessels in the brain parenchyma) (27). One day post-infection (dpi), sterile saline (untreated), minocycline (0.5 mg/kg/d; an inhibitor of microglia activation), PEA (0.5 mg/kg/d), AmpB (3 mg/kg/d), or a combination of AmpB (3 mg/kg/d) + PEA (0.5 mg/kg/d) were administered via intraperitoneal injection (i.p.) in different groups of mice followed by every other day treatment for 28-dpi. The survivability of the untreated and treated *Cn*-infected mice was closely monitored throughout the duration of the protocol (Fig. 1B). On average but not statistically significant, PEA-treated mice that developed neurocryptococcosis survived longer (median: 13-dpi) than those in the untreated (median: 11-dpi) and minocycline-treated (median: 10-dpi) groups. Similarly, AmpB-treated animals showed much longer survival trend (median: 18-dpi) than PEA-treated mice, although this trend was not statistically significant. Interestingly, *Cn*-infected mice treated with combination of AmpB and PEA survived the longest (median: 26-dpi) with 3 out of 7 mice (approximately 42.9%) reaching to the end of the experiment (28-dpi; vs. untreated, *P* < 0.001; minocycline, *P* < 0.003; AmpB, not significant or ns; PEA, *P* < 0.002). In this regard, only one mouse in the AmpB group survived the whole experiment whereas all the untreated, minocycline-treated, and PEA-treated animals died by 15-, 24-, and 18-dpi, respectively. Clinical observations resulting from *Cn* infection included respiratory distress, neurological impairment, seizures, loss of mobility, and particularly, weight loss, which was a clear indicator of disease progression and mortality (Fig. 1C). These results indicate that while PEA alone had limited effect in extending the life of mice that developed cerebral cryptococcosis, when combined with AmpB, PEA showed a considerable additive effect.

**Fig. 1.**
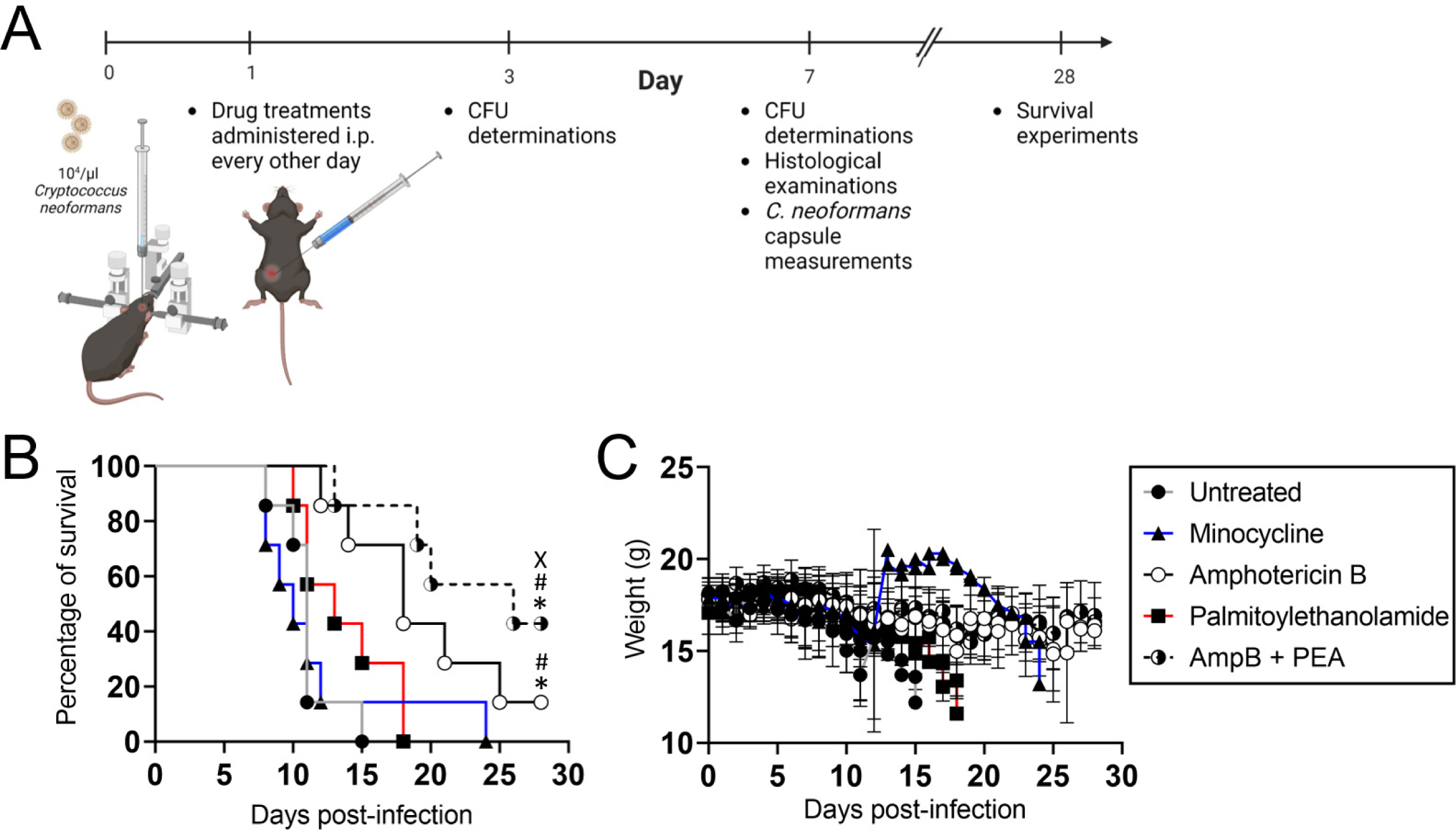
Palmitoylethanolamide (PEA) treatment prolongs survival of C57BL/6 mice infected with *C. neoformans* (*Cn*) when combined with Amphotericin B (AmpB). **(A)** Experimental timeline for the intracerebral (i.c.) *Cn* infection and every other day intraperitoneal (i.p.) treatment model used in this study. Mice (*n* = 7 animals per group) were infected with 10^4^ *Cn* strain H99 and administered treatments (e.g., saline (untreated), minocycline, AmpB, PEA, and AmpB + PEA) by i.p. injection every other day. Then, survival studies and colony forming units (CFU) determinations, histopathology, and microscopy were performed 3- and 7-days post-infection (dpi). The diagram was created with BioRender.com by Melissa E. Munzen. **(B)** Survival differences of C57BL/6 mice i.c. infected. Significance (*P* < 0.05) was calculated by log-rank (Mantel-Cox) analysis. *, #, and X indicate statistically different than untreated, minocycline-, and PEA-treatment groups, respectively. **(C)** Body weight was monitored for changes and development of clinical symptoms indicative of mice nearing endpoint. Each time point corresponds to mean weight and error bars denote standard deviations (SDs).

### AmpB and PEA combination significantly reduces fungal burden in *Cn*-infected brains

Coronal brain tissue sections were stained with Periodic acid-Schiff to identify the morphology of *Cn* in the host tissue during infection (Fig. 2). Untreated *Cn*-infected brains exhibit red-stained cryptococci (arrows) in the ventricular and the caudate nucleus (Fig. 2A). Minocycline- and PEA-treated brains displayed yeast cells accumulated in the margin of the cryptococcoma adhering to striatal tissue. Notably, brain tissue excised from mice treated with AmpB show minimal fungal colonization. *Cn*-infected mice treated with combination of PEA and AmpB displayed fungal cells in the lumen of the lateral ventricle. Mice i.c.-challenged with *Cn* [*n* = 6 mice per group; 6 colony forming unit (CFU) plates per animal were plotted] and treated with saline (untreated), minocycline, AmpB, PEA, or combination AmpB + PEA were evaluated for brain fungal load (Fig. 2B). There were no differences between the groups in fungal burden at 3-dpi. However, AmpB (mean: 3.72 x 10^6^ CFU/g tissue)- and AmpB + PEA (mean: 1 CFU/g tissue)-treated brains showed significantly lower fungal burden than untreated (9.14 x 10^6^ CFU/g tissue; *P* < 0.0001) or minocycline (8.04 x 10^6^ CFU/g tissue; *P* < 0.0001)-treated brain tissue at 7-dpi. The AmpB + PEA combination brains exhibited the lowest CFU number reduction of all the groups at the chosen dilutions. There were no differences in fungal burden between untreated, minocycline-and PEA-treated mice. Overall, our findings indicate that PEA is not fungicidal, although it enhances cryptococcal killing when combined with AmpB.

**Fig. 2.**
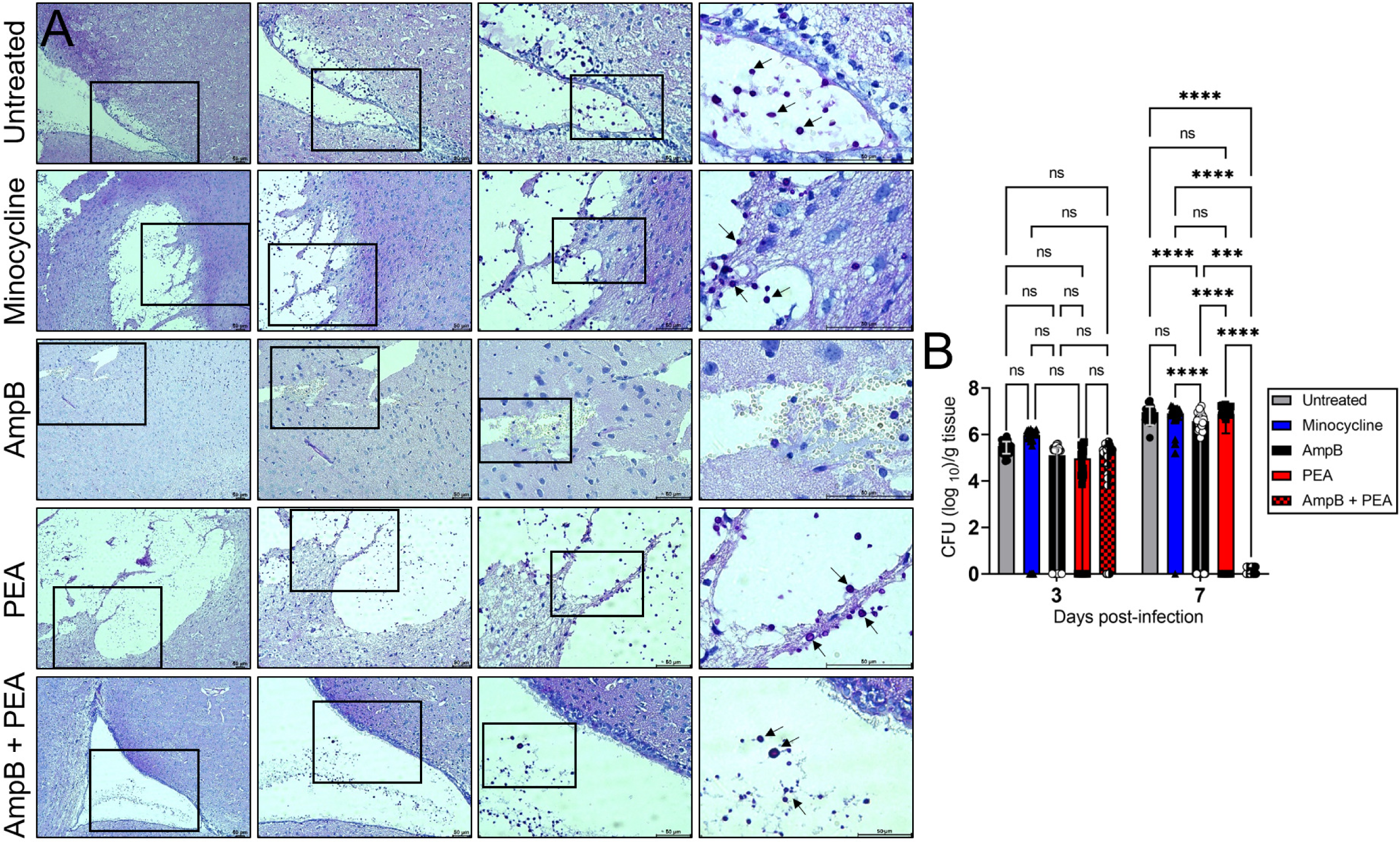
PEA and AmpB combination reduces cryptococcal burden in brains of mice 7 days post-infection (dpi). **(A)** Periodic acid-Schiff-stained 4 μm brain sections indicating infection by *Cn* in C57BL/6 mice (*n* = 3 mice per group). Mouse brains were excised 7-dpi. Representative 10X (left panel), 20X (left center panel), 40X (right center panel), and 100X (right panel) magnifications (red-stained *Cn* cell wall; scale bar: 50 μm) are shown. Panel images are a magnification of the black rectangle in the corresponding left-stained section to display tissue morphology surrounding cryptococcoma in each treatment group. Arrows indicate cryptococci. **(B)** Fungal burden (CFU) in brains collected from *Cn* H99-infected mice (*n* = 6 mice per group) at 3- and 7-dpi. Quantification of viable yeast cells from infected animals were determined by CFU counting from two dilutions per mouse (*n* = 6 plates per animal) in phosphate-buffered saline (PBS). CFU determinations are based on detectable colonies at the defined concentrations in PBS (for 3-dpi, 1:1,000 and 1:10,000; for 7-dpi, 1: 10,000 and 1:50,000). Each symbol represents a single CFU determination (*n* = 36 plates per group). Bars and error bars denote means and SDs, respectively. Significance (****, *P* < 0.0001; ***, *P* < 0.001) was calculated by one-way analysis of variance (ANOVA) and adjusted using Tukey’s *post hoc* analysis. ns denotes comparisons which are not statistically significant.

### AmpB and PEA combination stimulate strong glial cell responses around the brain region of infection

We have previously shown that *Cn* forms cryptococcomas in mouse striatal tissue after i.c. infection, mimicking CME in humans, particularly those cases with brain parenchyma involvement (28). Thus, we examined morphological changes in all saline (untreated), minocycline, AmpB, PEA, or AmpB + PEA treated *Cn*-infected mice. Brain tissue was processed for histology and sections were stained with hematoxylin & eosin, which allows us to observe tissue morphology. All treatment groups display encephalomalacia or cyptococcoma formation in the striatum (Fig. 3). Untreated brain tissue showed extensive necrosis that expanded throughout the basal ganglia (e.g., striatum, globus pallidus, and caudate nucleus; Fig. 3, top panels). Microgliosis and *Cn*-induced necrosis (red arrow) were also evident around the border of the cryptococcoma. In the minocycline-treated brains, extensive encephalomalacia that extended to the overlying cerebral cortex was observed (Fig 3, second panel row from the top). In addition, *Cn* invaded the tissue surrounding the cryptococcoma causing ischemic neuronal necrosis (blue arrow) and weak microglial response (red arrow), which is characteristic of GXM accumulation and the direct negative effect of minocycline in these glial cells (25). Interestingly, tissue from AmpB-treated mice evinced marked microgliosis (blue arrow) and astrocytic response (red arrow) in tissue adjacent to the cryptococcoma or encephalomalacia (middle row panels), while tissue from PEA-treated mice displayed noticeable microgliosis (red arrow) and evident swollen endothelial cells (blue arrow). Lastly, *Cn*-infected mice and treated with combination of AmpB + PEA exhibited microgliosis (red arrow) around the margin of the cryptococcoma (Fig. 3, lower panels). In summary, analysis of our histological data demonstrate that AmpB + PEA stimulate robust glial cell responses that may be critical to combat *Cn* brain infection.

**Fig. 3.**
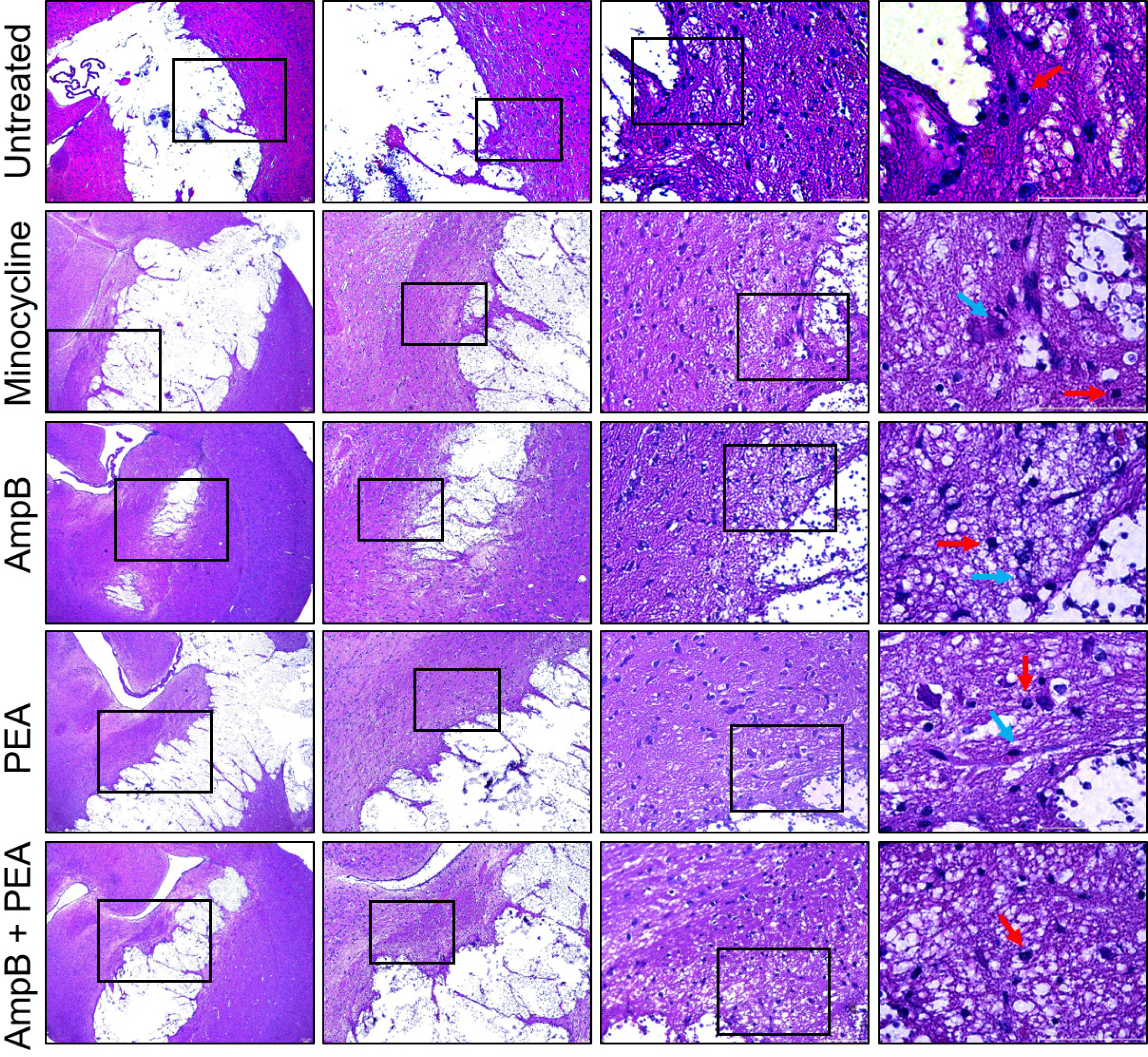
Robust glial cell responses were observed around the brain region of infection of mice treated with AmpB and PEA combination. Hematoxylin & eosin (7-dpi)-stained brain sections (4 μm thickness) from i.c. infected mice with *Cn* H99 (*n* = 3 per group) and treated with saline (untreated), minocycline, AmpB, PEA, or AmpB + PEA. Representative 10X (left panel), 20X (left center panel), 40X (right center panel), and 100X (right panel) magnifications images are shown. Arrows indicate histological changes described in the result section text. Scale bars: 50 μm.

### AmpB and PEA combination reduces *Cn* GXM released in brain tissue

*Cn* GXM is abundantly secreted during infection resulting in immunosuppressive effects for the host. Hence, we examined the impact of these various drug treatments on GXM accumulation in tissue during cerebral cryptococcosis (Fig. 4). GXM was red-pink stained in brain tissue using the specific monoclonal antibody (mAb) 18B7 (Fig. 4A). The GXM intensity in tissue sections was analyzed using NIH Image J color deconvolution tool software. Immunolabeling of untreated *Cn*-infected brain tissue evinced extensive GXM accumulation that diffused inside the lateral ventricle. Minocycline-injected animals displayed GXM concentrated in the area of encephalomalacia that spread to surrounding brain tissue. In contrast, *Cn*-infected brains from PEA treated mice showed localized polysaccharide accumulation inside the cryptococcoma without dissemination to surrounding tissue (Fig. 4A, middle panels). Interestingly, AmpB-treated groups (either alone or in combination with PEA) demonstrated a significant reduction in GXM accumulation in brain tissue (Fig. 4A and B). Further, brains from mice treated with AmpB + PEA combination showed positive immunostaining exclusively in plug-like structures (arrow) formed and adhered to ependymal cells in the lateral ventricle, without brain parenchyma involvement (Fig. 4A, lower panels). Analysis of the intensity of GXM staining in brain tissue images from *Cn*-infected mice after the various drug treatments revealed that AmpB alone and in combination with PEA had significantly lower GXM intensity or accumulation in tissues than minocycline-(*P* < 0.05) and PEA-treated (*P* < 0.01) groups (Fig. 4B). Our data show that AmpB regulates fungal capsular polysaccharide release, which may have important implications for impacting the progression of neurocryptococcosis.

**Fig. 4.**
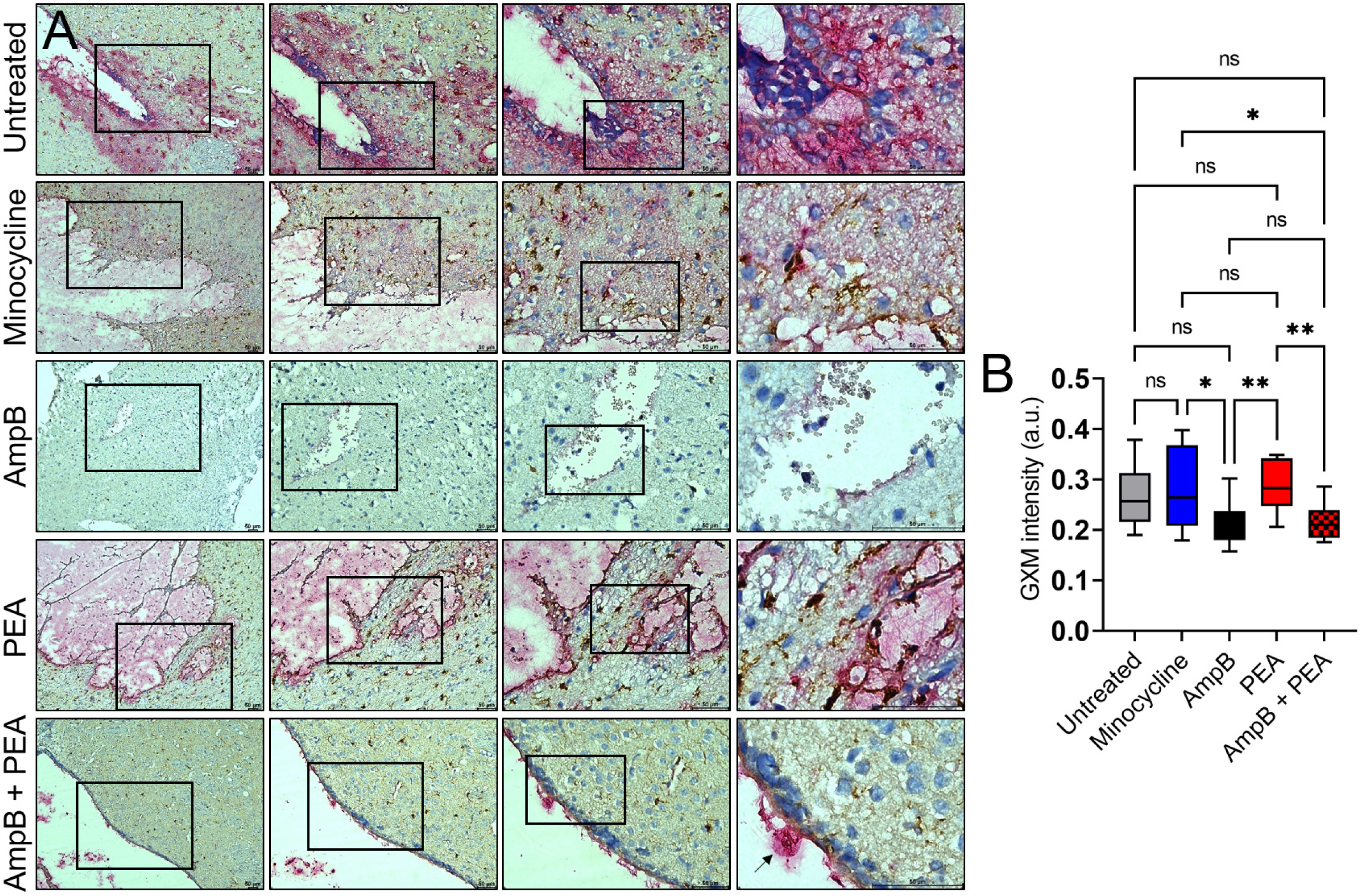
C57BL/6 mice treated with AmpB or AmpB + PEA showed significant reduction in *Cn* glucuronoxylomannan (GXM) secretion. **(A)** Representative images of brain tissue sections (7-dpi) from *Cn* H99-infected mice treated with saline (untreated), minocycline, AmpB, PEA, or AmpB + PEA co-stained with GXM-specific monoclonal antibody (mAb 18B7; red-pink) and ionized calcium binding adaptor molecule-1 (Iba-1; brown) marker for microglia. Representative 10X (left panel), 20X (left center panel), 40X (right center panel), and 100X (right panel) magnifications are shown. Panel images are a magnification of the black rectangle in the corresponding left-stained section to display tissue morphology surrounding cryptococcoma in each treatment group. Arrow indicates plugs adhered to the ependymal cells. Scale bars: 50 μm. **(B)** Quantification of GXM intensity. Regions of GXM release were measured (*n* = 15 fields per group). Boxes and whiskers denote means and SDs, respectively. Significance (**, *P* < 0.01; *, *P* < 0.05) was calculated by one-way ANOVA and adjusted using Tukey’s *post hoc* analysis. ns denotes comparisons which are not statistically significant.

### PEA has no effect in reducing *Cn* capsular volume *in vivo*

The capsule of *Cn* is the primary virulence factor of the fungus and is incredibly dynamic and responsive to changes in the microenvironment. Large capsule volume is correlated with a robust inflammatory response and respiratory distress (29). Also, capsular enlargement reduces phagocytosis and enhances *Cn* survival during infection (30). To validate our results indicating that AmpB treatment reduces *Cn* GXM released in brain tissue, we assessed the impact of the different drug treatments on capsular size and volume in brain tissue homogenates using India ink staining and light microscopy (Fig. 5A). Quantification of microscopic images (*n* = 3 mice and ≥ 200 cells per group; Fig. 5B) confirmed that treatment with AmpB (1,208.2 μm^3^; *P* < 0.05) and AmpB + PEA (1,259.9 μm^3^; *P* < 0.05) significantly reduced *Cn* capsular volume compared to minocycline treatment (1,643.2 μm^3^). Surprisingly, no differences in capsular volume were observed between untreated (1,394.6 μm^3^), PEA (1,350.1 μm^3^), AmpB (1,208.2 μm^3^), and AmpB + PEA (1,259.9 μm^3^). Our results suggest that AmpB, on average, has a negative effect on cryptococcal capsular production and release in brain tissue, thus, facilitating fungal clearance during infection.

**Fig. 5.**
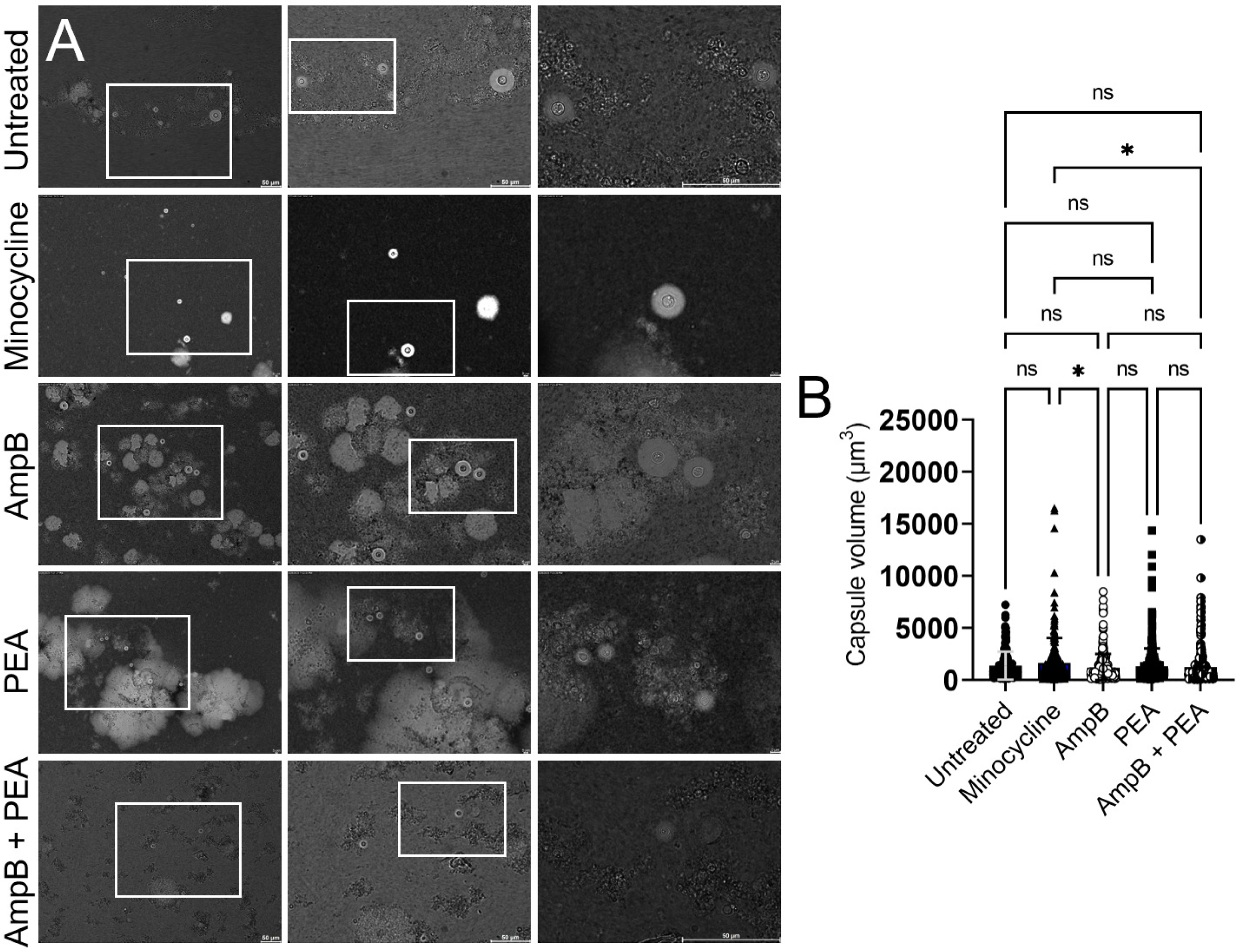
Brains from C57BL/6 mice treated with AmpB or AmpB + PEA exhibit cryptococci with considerable capsule size decrease. **(A)** Images of brain homogenates (7-dpi) from mice i.c. infected with 10^4^ *Cn* cells and treated with saline (untreated), minocycline, AmpB, PEA, or AmpB + PEA. Fungal cells were stained with India Ink. Each image was examined by light microscopy using a Leica DMi8 inverted microscope and images captured with a Leica DFC7000 digital camera using LAS X digital imaging software. **(B)** Capsule volume (V = 4/3 π(R^2^-r^2^) for *Cn* cells in brain homogenates from each group was calculated using Leica LAS X software. Brain homogenates from 3 mice per group were analyzed, and ≥ 200 cells were measured. Bars and error bars denote means and SDs, respectively. Significance (*, *P* < 0.05) was calculated by one-way ANOVA and adjusted using Tukey’s *post hoc* analysis. ns denotes comparisons which are not statistically significant.

### PEA causes similar microglia morphological changes than minocycline during *Cn* brain infection

Microglia are the principal immune cell in the brain responsible for phagocytosis, immune cell recruitment, and communication via cytokine signaling (31). We recently reported that microglia can take diverse morphological phenotypes after *Cn* i.c.-infection, ramified, activated, phagocytic/amoeboid, dystrophic, or rod-shaped, which may be associated with neurocryptococcosis progression and outcome (28). Therefore, we monitored the impact of each drug treatment on the phenotypic landscape of microglia during cryptococcal infection (Fig. 6). Microglia and GXM were immunolabeled using ionized calcium binding adaptor molecule-1 (Iba-1; brown)- and 18B7 (pink-red)-binding mAbs, respectively (Fig. 6A). Images of each group treatment were visualized under the microscope and the microglia morphology documented as previously described (25). *Cn*-infected brains from untreated mice presented the following microglia phenotype distribution: activated (38.52%) > ramified (31.03%) > rod-shaped (15.46%) > phagocytic (11.59%) > dystrophic (3.4%) (Fig. 6B). Although minocycline treatment inhibited microglial activation (32.46%), this drug resulted in the highest percentage of phagocytic (42.8%) and dystrophic (9.58%) microglia and the lowest percentage of ramified (6.03%) microglia. In addition, brains from minocycline-treated mice had the second lowest percentage of rod-shaped (9.12%) microglia after AmpB-treated (8.46%) mice. Furthermore, AmpB-treatment resulted in the highest percentage of ramified (44.58%) microglia and the lowest percentage of activated (22.78%) microglia. Tissue from AmpB-treated mice also had 20.93% and 3.25% of phagocytic and dystrophic microglia, respectively. The phenotypic profile of microglia in PEA-treated tissue was similar to those observed in the minocycline group including 39.32% phagocytic, 35.77% activated, 11.69% rod-shaped, 10.37% ramified, and 2.85% dystrophic morphology. Moreover, AmpB + PEA-treated animals evinced similar microglia morphological patterns than untreated mice including 32.48% activated, 30.18% ramified, 14.86% rod-shaped, 20.18% phagocytic, and 2.29% dystrophic cells. Our findings suggest that *Cn* brain infection and different drug treatments causes phenotypic microglial alterations, which may have important consequences in their responses to fungal infection and clearance.

**Fig. 6.**
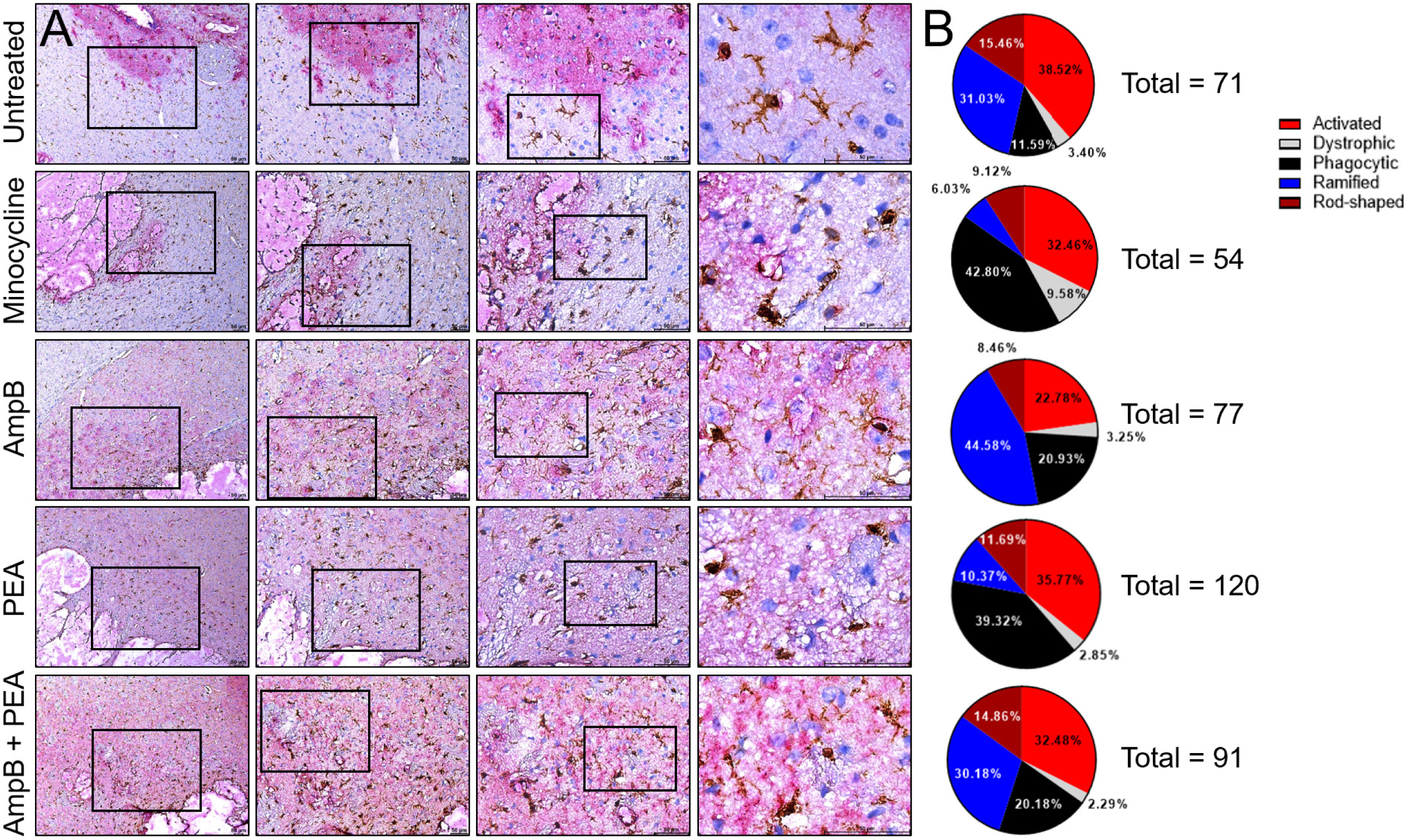
Differential microglial morphology in brain tissue infected with *Cn* after treatment with AmpB, PEA, or combination. **(A)** Representative images of brain tissue sections (7-dpi) from *Cn* H99-infected mice (*n* = 3 mice per group) treated with saline (untreated), minocycline, AmpB, PEA, or AmpB + PEA co-stained with mAb 18B7 (GXM, red-pink) and Iba-1 (microglia, brown). Representative 10X (left panel), 20X (left center panel), 40X (right center panel), and 100X (right panel) magnifications are shown. Panel images are a magnification of the black rectangle in the corresponding left-stained section to display microglia morphological changes near a cryptococcoma in each treatment group. Scale bars: 50 μm. **(B)** Pie charts showing the percentage of microglial morphology distribution (e.g., activated, ramified, dystrophic, phagocytic/amoeboid, and rod-shaped cells) in brain tissue during infection with *Cn* H99 and different treatments. Microglial phenotype abundance was visualized using light microscopy and classified according to their morphology.

### PEA and AmpB combination prolong i.c.-infected mouse survival, but with less efficacy to current standard of treatment

Since AmpB and 5-FC are standard of treatment for individuals with CME, we compared the efficacy of PEA and AmpB combination and AmpB and 5-FC combination in extending *Cn-*infected mice survival (Fig. 7). On average, *Cn*-infected and AmpB + PEA-treated mice exhibited higher survivability (median: 20-dpi) relative to untreated (median: 12-dpi; *P* < 0.03) and PEA + 5-FC (median: 14-dpi; trend but ns) animals (Fig. 7A). All untreated mice and PEA + 5-FC-treated groups died by 20-dpi. On the contrary, all mice infected with *Cn* and treated with AmpB + 5-FC survived (100%; vs. Untreated, *P* < 0.001; PEA + 5-FC, *P* < 0.001) whereas only 33% (2 out of 6) of the animals in the AmpB + PEA group (*P* < 0.02) survived for 28-dpi when the experiment was finalized. Body weight was monitored and was also a clear indicator of disease progression, with mortality being associated with 25% weight loss (Fig. 7B). These results show that while the combination of AmpB and PEA has positive effects in prolonging the life of mice with cerebral cryptococcosis, this effect is limited when compared with the efficacy of AmpB and 5-FC, the current standard of patient care.

**Fig. 7.**
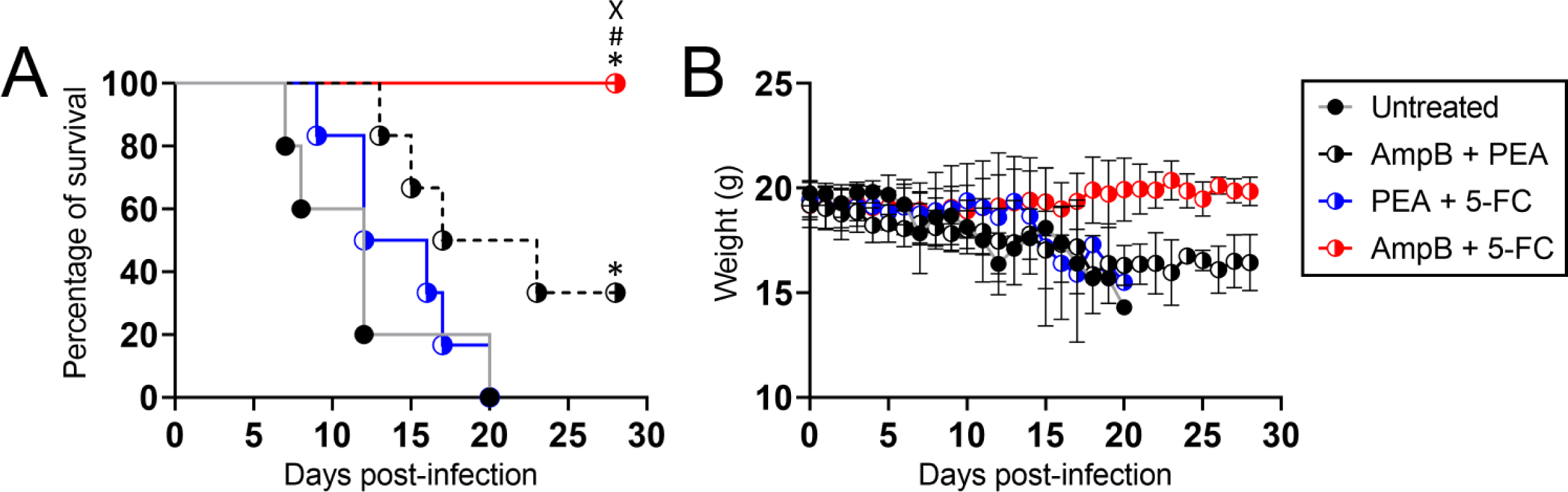
Combination of AmpB and PEA prolongs survival of C57BL/6 mice, although not with efficacy comparable to AmpB and 5-Fluorocytosine (5-FC). **(A)** Survival differences of C57BL/6 mice i.c. infected with 10^4^ *Cn* strain H99 (*n* = 6 per group) and every other day given i.p. treatment with saline (untreated), AmpB + PEA, PEA + 5-FC, or AmpB + 5-FC. Significance (*P* < 0.05) was calculated by log-rank (Mantel-Cox) analysis. *, #, and X indicate statistically different than untreated, AmpB + PEA-, and PEA + 5-FC-treatment groups, respectively. **(B)** Body weight was monitored for changes and development of clinical symptoms indicative of mice nearing endpoint. Each time point corresponds to mean weight and error bars denote SDs.

## DISCUSSION

In this study, we demonstrated that PEA has limited effect in prolonging murine survivability upon direct brain infection with *Cn*. PEA is an immunomodulatory molecule, and its effects in extending mouse survival are related to the high activation of microglia in brain tissue that may aid in combating *Cn* infection. Our data show that 75% of microglia during *Cn* brain infection had an activated or phagocytic phenotype, which is critical for microbial clearance (32). For example, PEA increases microglia-mediated phagocytosis of bacteria without inducing considerable inflammation (33), prevents rheumatic fever in susceptible children (34), and alters the course of influenza infection (24). However, its limitation in controlling *Cn* infection is because PEA has no direct effect in killing the fungus, reducing cryptococcoma formation, or decreasing fungal capsular volume and GXM secretion. Interestingly, PEA provides a substantial additive effect in combating cerebral cryptococcosis when combined with AmpB, which although toxic to the host, is a potent anti-cryptococcal agent, validating the critical importance of antimicrobial compounds in stimulating the host immunity. Combination of AmpB and PEA significantly and surprisingly decreased CFU numbers in *Cn* brain infected mice relative to all the other tested groups. Despite cryptococcoma formation in brain tissue of animals treated with AmpB and PEA combination, histopathology shows considerable microgliosis surrounding the brain-lesions, which is consistent with fungal clearance. Although i.c. infected mice treated with either AmpB or PEA elicited a strong glial response, it is plausible to consider that the AmpB and PEA combination provides an ideal balance between the potency of the antifungal drug and the lipid in killing the fungus. It is possible that PEA, by being a hydrophobic molecule, binds to the cell membrane ergosterol and serves as a vehicle in facilitating AmpB-mediated pore formation, killing the cell (35). These hypotheses are possible given that untreated and minocycline-treated brains showed moderate to weak glial responses and the untreated tissue exhibited ischemic necrosis, a manifestation reported in patients with CME (36, 37) and in this model of infection (25).

Untreated and minocycline- or PEA-treated *Cn* infected brains exhibited extensive accumulation of GXM in tissue. Additionally, cryptococci isolated from the brains of these groups showed large capsule size and volume relative to the AmpB-treated groups of animals. Enlargement of the capsule reduces phagocytosis during infection and dramatically enhances virulence through oxidative stress resistance (30, 38). However, cryptococcal GXM accumulation showed different patterns with untreated and minocycline-treated tissue displaying polysaccharide intensity throughout tissue surrounding the cryptococcoma, whereas PEA-treated tissue exhibited higher GXM intensity inside of cryptococcomas or region of encephalomalacia. It is conceivable that differential glial cell responses (e.g., microglia and astrocytes) between these groups modulate the extension (e.g., untreated and minocycline) of GXM secretion from or containment (e.g., PEA) within the cryptococcoma. This is important because GXM inhibits T cell proliferation and reduces leukocyte recruitment needed for fungal clearance (39). Notably, AmpB and AmpB + PEA brains infected with the fungus evinced little or localized tissue secretion of GXM, respectively, which is likely due to the smaller capsular volume shown by the cryptococci and reduction in proliferation or killing of the fungi. AmpB reduces capsule size and serum polysaccharide, a described mechanism for this antifungal drug efficacy in cryptococcosis (40). Regardless of its high toxicity, AmpB has been used for the treatment of disseminated cryptococcal disease for over 70 years (41), and novel anti-cryptococcal compounds causing minimal toxicity are urgently needed.

Microglia are dynamic resident immune cells of the central nervous system that can modify their phenotype based on their responses to various pathological insults (42). Microglia engulf cryptococci, which can survive and replicate inside these cells (43), avoiding immune recognition. The morphological distribution of microglia in brain tissue after *Cn* brain infection and the various treatments differed among the groups and may have important predictor implications in reducing mortality. For instance, untreated *Cn*-infected brains showed the highest and lowest percentages of rod-shaped and phagocytic morphologies by microglia, respectively. Descriptions of rod-shaped microglia, which present elongated nuclei, few processes, and are most notable in chronic neurological disorders, are scarce. The function of these cells remains under investigation, but it is believed that they may provide neuroprotection and trophic support (44, 45). Hypertrophic microglia are thought to be actively responding to injury (46, 47) and were most abundant in untreated *Cn*-infected brains mostly due to tissue destruction related to encephalomalacia. Hypertrophic microglial densities are directly related to cortical atrophy in Alzheimer’s disease and inversely related to high density of functional neurons in tissue (48). Thus, accumulation of activated microglia may occur in damaged blood vessels or microinfarctions during cryptococcal infection. It is also possible that GXM accumulation makes activated microglia anergic or unresponsive, thus, the low phagocytic or amoeboid morphology displayed, which may facilitate the proliferation and survival of the fungus, enhancing the progression of the cerebral disease. Both minocycline- and PEA-treated tissues presented approximately three quarters of all microglial cells activated and phagocytic, but animals in both groups have low number of ramified microglia. Activated or hypertrophic microglia has been linked to oxidative stress and inflammation in neurological diseases such Alzheimer’s (49), Parkinson’s (50), stroke (51), and others. Moreover, minocycline-treated animals exhibited the highest number of dystrophic microglia, which have fragmented processes, probably due to cell dysfunction. Dystrophic microglia are found in high numbers in vulnerable individuals who suffer from brain degeneration (52, 53) or senescence (44) due to aging or neurological diseases. Interestingly, brains excised from AmpB and combination of AmpB and PEA groups showed similarly balanced activated, phagocytic, ramified microglia distribution percentages. It is conceivable that the highest number of ramified and rod-shaped microglia present in AmpB- and combination-treated animals, respectively, makes a difference in the fungal persistence or clearance observed in these groups, possibilities that can be tested in future studies. Elucidating the impact of microglial responses or phenotypes in brain tissue infected with *Cn* is necessary given their central role in many neurodegenerative diseases, including CME, which leads to cognitive decline and mortality in patients (54).

Treatment of *Cn*-infected mice with combination of AmpB + PEA revealed less efficacy than the recommended standard combination of AmpB + 5-FC. All the mice treated with the standard of care combination survived through the duration (28-dpi) of the experiment. However, mortality associated with CME in humans even with optimal treatment remains high (55). Therefore, novel treatments that rapidly improve the clinical manifestation associated with CME such as intracranial pressure and high fungal burden in the cerebrospinal fluid. Here, we demonstrated that PEA is not an effective antifungal drug but may be beneficial in modulating the immune response in combination with an effective fungicidal drug. Further pre-clinical studies investigating the efficacy of immunomodulator molecules for the treatment of CME could provide medical professionals with additional cost-effective and safe antifungal therapy options of care especially in developing countries.

## MATERIALS AND METHODS

### Ethics statement

All animal studies were conducted according to the experimental practices and standards approved by the Institutional Animal Care and Use Committee (IACUC) at the University of Florida (Protocol #: 202011067). The IACUC at the University of Florida approved this study.

### Cn

*Cn* strain H99 (serotype A) was isolated and kindly provided by John Perfect at Duke University. Yeasts were grown in Sabouraud dextrose broth (pH 5.6; BD Difco) for 24 h at 30°C in an orbital shaker (Thermo Fisher) set at 150 rpm (to early stationary phase).

### i.c. infection with *Cn*

C57BL/6 female mice (6-8 weeks old; Envigo) were anesthetized using isoflurane (3–5% for induction and 1–2% maintenance; model: VetFlo Vaporizer Single Channel Anesthesia System, Kent Scientific), placed in prone position over a heating pad (model: RightTemp Jr., Kent Scientific), and prepped using standard aseptic techniques. A local anesthetic, bupivacaine or ropivacaine (0.05%; Covetrus), was administered subcutaneously in the incision. The fur on the skull was carefully shaved off and the animal was securely placed in a stereotaxic apparatus (model: 940; Kopf Instruments).

Using a small hand-held microdrill (model: Ideal microdrill; Braintree Scientific), the skull was thinned until the underlying dura mater was visible and a 26 G Hamilton syringe was brought to the correct stereotaxic position and lowered until it touched the exposed dura. The craniotomy was around 1 mm in diameter and the correct brain coordinates were identified using a stereotaxic brain atlas (e.g., The Allen Mouse Brain Atlas; https://mouse.brain-map.org/static/atlas). Based on our recent study (25) and others (56, 57) using the *Cn* H99 strain, a 1-μL suspension containing 10^4^ cryptococci in sterile saline was injected into the striatum [Stereotaxic coordinates: x (medial/lateral), -2; y (anterior/posterior), 0.2; z (dorsal/ventral), - 3.5)] using a 26 G Hamilton syringe connected to a pump (model: UltraMicroPump3; World Precision Instruments). We injected the fungal inoculum in a 1-μL volume to avoid tissue damage or diffusion of the cryptococci to other regions of the brain. The skin incision on the dorsal head was closed with sterile nylon suture and 2-4% topical chlorhexidine solution was be applied over the closed incision. After the surgery, mice were placed on a clean recovery cage.

### Administration of drug treatments

One-dpi, mice were injected with a 100 μL solution of sterile saline, minocycline (0.5 mg/kg/day (d); diluted in sterile saline; ACROS Organics), AmpB (3 mg/kg/d; diluted in sterile saline; AmpB; Gibco), PEA (0.5 mg/kg/d; diluted in sterile saline; Tokyo Chemical Industry), or AmpB (3 mg/kg/d) + PEA (0.5 mg/kg/d) combination i.p. every other day and monitored for survivability. In separate infections, mice were euthanized at determined time points via CO_2_ inhalation and brain tissues were excised for processing for determination of CFU numbers and histopathological studies. Similarly, 5-fluorocytosine (100 mg/kg/d; diluted in sterile saline; 5-FC; ThermoFisher) was used in combination with PEA (0.5 mg/kg/d) or AmpB (3 mg/kg/d) and survivability was determined. The survival end points were inactivity, tachypnea, or loss of ≥ 25% of body weight from baseline weight. We monitored the mice twice daily for clinical signs, dehydration, and weight loss. Animals showing signs of dehydration or that lost more than 10% weight received supportive care such as 1 mL of parenteral fluid supplementation (saline) and moist chow on the cage floor was provided.

### CFU determinations

Brains were excised from euthanized mice and weighed 3- and 7-dpi. The brain tissue was homogenized in 5 mL of sterile phosphate buffered saline, serially diluted, a 100 μL suspension was plated on Sabouraud dextrose agar (BD Difco) and incubated at 30°C for 48 h. Quantification of viable yeast cells from infected animals were determined by CFU counting of two dilutions per animal (*n* = 6 per day).

### Brain histology

The brains were harvested and immersed in 4% paraformaldehyde (Fisher) overnight. Then, brains were washed 3X with sterile saline for 1 h, embedded in paraffin, 4 μm coronal sections were serially cut using a cryostat (Tanner Scientific, model: TN50), fixed onto glass slides, and subjected to hematoxylin & eosin or Periodic acid-Schiff staining to examine tissue or fungal morphology, respectively. GXM (MAb 18B7 is an anti-cryptococcal GXM IgG1 generated and generously provided by Arturo Casadevall at the Johns Hopkins Bloomberg School of Public Health; 1:1,000 dilution) and Iba-1 (rabbit anti-human Iba-1; 1:1,000 dilution; FujiFilm Wako) specific Ab (conjugated to horseradish peroxidase; dilution: 1:1,000; Santa Cruz Biotechnology) immunostaining to assess capsular release and distribution and microglial phenotype, respectively, near cryptococcomas. The slides were visualized using a Leica DMi8 inverted microscope, and images were captured with a Leica DFC7000 digital camera using LAS X digital imaging software. GXM distribution in tissue sections at 10X magnification (*n* = 15 fields per brain) was calculated using NIH Image J color deconvolution tool software (version 1.53q). The mean color intensity of the GXM for each treatment group was plotted in Prism version 9.5 (GraphPad). The images were examined and analyzed by Dr. Mohamed F. Hamed, a veterinary pathologist.

### Capsule measurements with India ink

The capsules of cryptococci present in the brain tissue of infected mice were measured from 10-μL of homogenates prepared in phosphate-buffered saline. India ink stain (BD Scientific) was diluted 1:5 in sterile milli-Q water and used to visualize the capsules under light microscopy. Images were taken with a Leica DMi8 inverted microscope and DFC7000 T digital camera. The diameters of both capsule and cell body were measured using the Leica software platform LAS X. Capsule volume was calculated for ≥ 200 cells per group (*n* = 3 animal per group) using the volume formula, where R = radius of capsule and r = radius of cell body: capsule volume (V) = 4/3 π(R^2^-r^2^).

### Statistical analysis

All data were subjected to statistical analysis using Prism 9.5 (GraphPad). Differences in survival rates were analyzed by the log-rank test (Mantel-Cox). *P* values for multiple comparisons were calculated by one-way analysis of variance (ANOVA) and were adjusted by use of the Tukey’s *post-hoc* analysis. *P* values of <0.05 were considered significant.

## FIGURE LEGENDS

## ACKNOWLEDGEMENTS

M.E.M., M.R-G, M.F.H., C.L.C-N, M.R.D., and L.R.M. were supported by the National Institute of Allergy and Infectious Diseases (NIAID award # R01AI145559) of the US National Institutes of Health (NIH). V.E. was supported by the UFCD’s Comprehensive Training Program in Oral Biology (NIH NIDCR Award # T90DE021990/R90DE022530). K.A. was funded by The Florida Department of Health Ed and Ethel Moore Alzheimer’s Disease Research Program (Award # 21A12). The funders had no role in the study design, data collection and analysis, decision to publish, or preparation of the manuscript.

## AUTHORSHIP CONTRIBUTIONS

All authors contributed to the project design and experimental procedures, analyzed data, provided the figure presentation, and manuscript writing.

